# Development of ultrasound molecular contrast agents by site-specific conjugation of gas vesicles to antibodies

**DOI:** 10.1101/2025.07.21.665993

**Authors:** Sydney Turner, Adree Bhattacharjee, Lu Diao, Min Zhao, Siyuan Zhang, Sangpil Yoon

## Abstract

Ultrasound technology is a powerful tool for medical imaging and diagnosis especially when used in conjunction with a contrast agent to target proteins of interest. However, current ultrasound contrast agents are lacking the combination of specificity, stability, and size optimization. In this study we introduce and confirm the feasibility of using nanometer sized gas vesicles (GV) chemically click conjugated to antibodies (mAb) to make the first site-specific mAb-GV conjugates as a durable cancer cell targeting ultrasound molecular contrast agents. Protein expression of human epidermal growth factor receptor 2 (HER2) and programmed death-ligand 1 (PD-L1) were tested along with the antibody targeting efficiency using cancer cell lines and primary cells isolated from tumor bearing mice. The mAb conjugation was optimized to a site-specific method using the mAb glycans and tested with the addition of a clinically used mAb to target trophoblast cell surface antigen 2 (Trop-2). The developed contrast agents utilize the stability of GVs and the specificity of antibodies to label cancer biomarkers in various types of tumors for ultrasound imaging.

## 1. Introduction

Cancer is currently the second leading cause of death in the United States with 1,958,310 new cancer cases and 609,820 cancer deaths in 2023.^[1]^ Ultrasound technology provides a noninvasive imaging option for diagnosing and monitoring, though it is limited in the ability to distinguish between tissue types and structure unless contrast agents are used. Established cancer treatments such as chemotherapy and radiation are rapid, effective, and can be used broadly, but due to the detrimental side effects, a new push for immunotherapies has arisen.^[2]^ Immunotherapy utilizes monoclonal antibodies (mAb) to suppress and kill cancer cells by labeling antigens on their surface, which then increases the stimulation of the immune system.^[3]^

Three mAbs were chosen for this study due to their impact and wide applicability. Human epidermal growth factor receptor-2 (HER2) is overexpressed in many types of solid tumors such as non-small-cell lung, colorectal, gastric, bladder, biliary tract, and gastroesophageal junction cancers, but is the most common in breast cancer leading to excessive cell proliferation.^[4]^ The blocking of the HER2 with an anti-HER2 mAb inhibits the cell growth and division by interfering with the signal transduction pathway.^[5-6]^ Programmed cell death ligand 1 (PD-L1) is also overexpressed in many cancer types including non-small-cell lung, bladder, renal cell, gastric, and breast.^[7]^ In a healthy system, PD-L1 bind to PD-1 expressing T cells to regulate T cell activity. However, when cancer cells overexpress PD-L1, it disguises them from the immune system, hence the rationale for using an anti-PD-L1 mAb to visualize the immune evasion.^[8-9]^ Trophoblast antigen 2 (Trop-2) is a cell-surface transmembrane glycoprotein with an intercellular calcium signal transducer and is overexpressed in a number of cancers including breast, colon, colorectal, endometrioid endometrial, gastric, glioma, ovarian, pancreatic, and prostate.^[10-12]^ Trop-2 has been clinically explored as a tumor-associated target for chemotherapy.^[13]^ Specifically, Sacituzumab Govitecan is a clinically used antibody drug conjugate (ADC), combining an anti-Trop-2 targeting mAb (Sacit) with a chemotherapy drug, SN-38, to target tumor cells with antibody-mediated specificity. ^[14-16]^

There are several ways to conjugate into an mAb, such as using the 8 accessible thiol groups to form stable thioether bonds or using the 10-15 readily available lysine amine groups with the N-hydroxysuccinimide (NHS) method.^[17-18]^ However, both of these options involve additions to both the light and heavy chains regions of the mAb. Conjugating the binding region of the light chain inhibits the targeting of the cell markers and when adding a large particle, a conjugation anywhere on the light chain may hinder the targeting. Thus, we pursued site-specific labeling of mAb by tagging enzyme-mediated N-azidoacetylgalactosamine (GalNAz) on glycans on the CH2 region of the heavy chain (site-specific GalNAz method).^[19-20]^

Most mAb conjugate additions, such as drugs or fluorescent labels, are not naturally equipped with the maleimide, NHS, or galactose needed to bind to the mAb. Thus, bioorthogonal click chemistry is a valuable addition, as it can rapidly bind two prepared proteins without the need for a catalyst.^[21-22]^ The Staudinger ligation uses an azide and a phosphine to create a covalent amide bond.^[23]^ The strain-promoted azide-alkyne cycloadditions (SPAAC) uses ring strain energy produced by combining an azide with an alkyne to create a 1,2,3-triazole link.^[24]^ By preparing an mAb using the GalNAz method, an azide is ready for a click reaction with a second protein prepared with either a phosphine or an alkyne.^[20]^

Contrast agents with the ability to target cancer cells will improve ultrasound imaging and diagnosis.^[25-28]^ The most common ultrasound contrast agents currently used in clinic are different types of microbubbles, which are synthetic gas filled lipid or protein shells.^[29]^ However, the instability and short half-life of microbubbles, due to diffusion, limit the effectiveness especially after the time taken to conjugate them.^[30]^ Nanobubbles have been proposed as an alternative to microbubbles, because the smaller size allows them to enter small blood vessels for increased permeability and retention.^[31-33]^ Nanobubbles also have increased stability due to smaller sizes with increased surface tension. ^[34-35]^ Phase-change nanodroplet (PCND) uses perfluorocarbon to change phase from a liquid to a gas bubble when exposed to laser irradiation.^[36-37]^ PCNDs can be stored for days, but the size can decrease significantly from day one to day two and this instability limits conjugation opportunities.^[38]^

Gas vesicles (GV) are gas filled protein shells, which are found and purified from aquatic bacteria. GVs are stable for over a year and have a significantly longer half-life.^[25, 39-40]^ Not only are GVs genetically editable, but also the protein shell provides many available amine groups to be conjugated with NHS modified biomolecules.^[41-44]^ The GV type chosen for this study, *Anabaena flos-aquae* (AnaGV), has an average GV size ranging from 100 nm to 500 nm and has a ribbed shell of GV proteins, gas vesicle protein A (GvpA) and C (GvpC).^[45]^ GVs increase the contrast of ultrasound images and will highlight targeted areas when conjugated efficiently.^[25, 46-47]^

In this study, we compared the random NHS-azide conjugation method with the site-specific GalNAz conjugation method and tested our first ever mAb GV click chemistry complex (mAb-GV) with ultrasound imaging after cancer cells targeting. Cell lines and primary cells from tumor bearing mice were measured for HER2 and PD-L1 surface protein expression. Antibody targeting was verified using fluorescent microscopy. The NHS-azide conjugation method was tested by targeting the mAb-GV complex on cells and imaging with ultrasound. After noting possible limitations of the NHS method, the site-specific GalNAz method was optimized and comparisons were conducted using hydrodynamic range, zeta potential, degree of labeling (DOL), and flow cytometry after cell targeting. Trop-2 testing was performed with cell expression, research antibody, and clinic antibody verification. The final test compared the two mAb-GV conjugation methods with ultrasound imaging using four cell lines and five different antibodies to determine the superior technique based on brightness of ultrasound images that correlates to the labeling of proteins by mAb-GV.

## 2. Results

### 2.1 Optical validation of targeting efficiency of mAb

Before targeting cells with a mAb-GV conjugate and imaging it with ultrasound, we first sought to confirm cellular expression and antibody targeting using fluorescence and chemiluminescence. SK-BR-3 and EO771 were tested with flow cytometry, immunofluorescence, and Western blot, all of which showed high expression of HER2 on SK-BR-3 cells and expression of PD-L1 on EO771 (**Figure 1**A,C,D). After cell verification, mAbs to be used for GV conjugation were tested for targeting efficiency. The mAbs were labeled with fluorescence using the site-specific GalNAz method, making anti-HER2 mAb-647 (HER2-647) for SK-BR-3 and anti-PD-L1 mAb-488 (PD-L1-488) for EO771.^[48]^ The fluorescent intensity was compared between cells targeted with IgG-647 control, and cells targeted with either HER2-647 or PD-L1-488, showing significance for both cell lines (Figure 1 E,F). To further test targeting efficiency of anti-HER2 and anti-PD-L1 mAbs, we used primary cells. Flow cytometry was used to measure the PD-L1 and HER2 expression on tumor and spleen cells from a tumor bearing mouse after dissociation and magnetic isolated for Cd11b. The spleen showed a significantly higher PD-L1 expression compared to HER2, but the tumor, while having a higher PD-L1 expression also had a moderate HER2 expression (Figure 1B).

**Figure 1.**
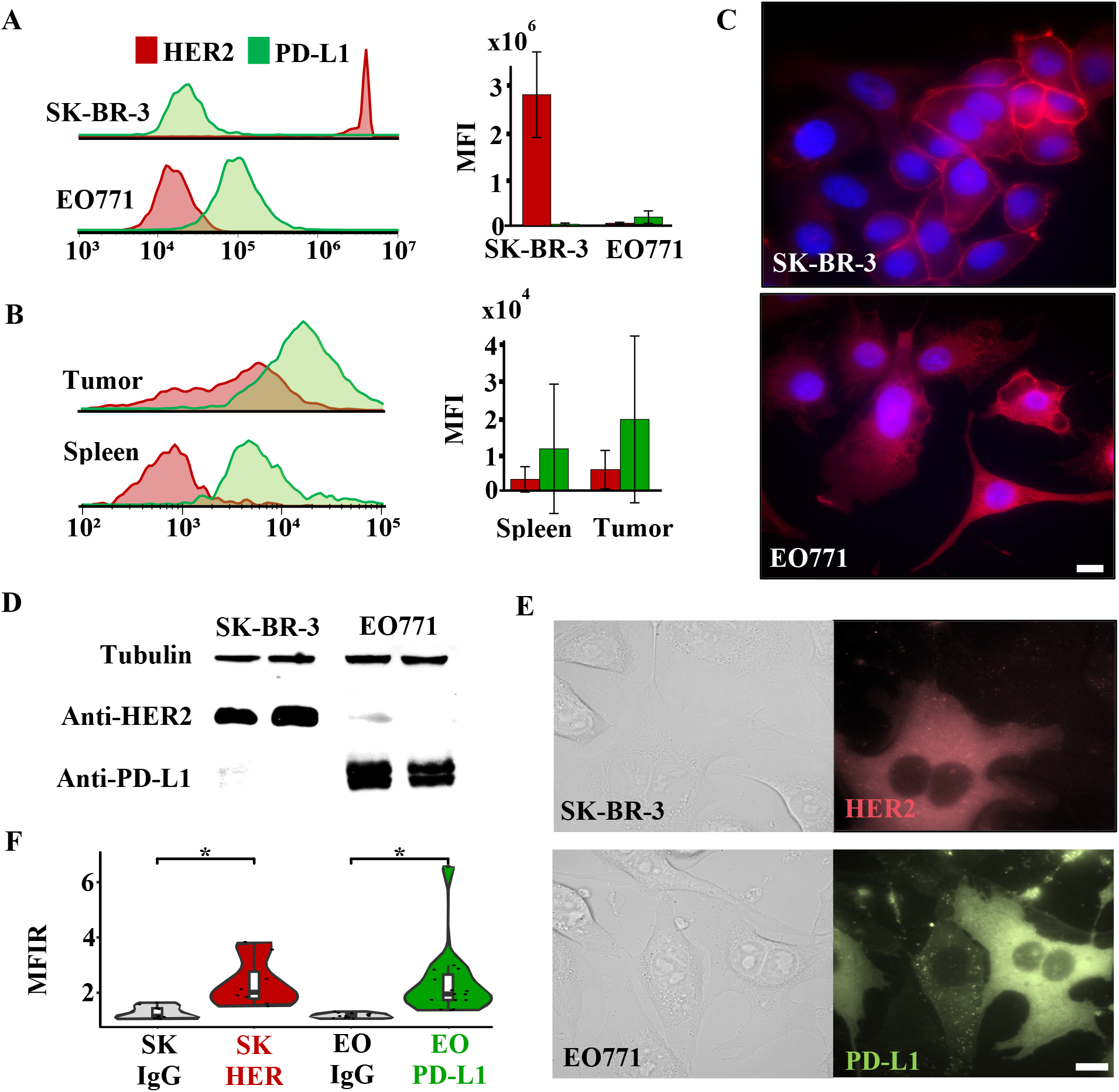
HER2 and PD-L1 cell expression and antibody targeting efficiency. A) Flow cytometry results of SK-BR-3 and EO771 targeted with anti-HER2 and anti-PD-L1 mAbs show stronger HER2 expression on SK-BR-3 than PD-L1 and stronger PD-L1 expression on EO771 than HER2. B) Flow cytometry of tumor and spleen primary cells taken from tumor bearing mice, isolated for CD11b, and targeted with anti-HER2 and anti-PD-L1 mAbs shows significant PD-L1 expression on both tumor and spleen cells, while HER2 expression is only significant in the tumor. C) Immunofluorescence (IF) test images show strong HER2 expression on SK-BR-3 and PD-L1 expression on EO771 labeled with Alexa 647 and cell nuclei labeled with DAPI. Scale bar = 10µm. D) Western blot results show stronger HER2 expression on SK-BR-3 than PD-L1 and stronger PD-L1 expression on EO771 than HER2 with Tubulin used as the control. E) mAbs labeled with fluorophores through the site-specific conjugation method were targeted to cells showing HER2 in Alexa 647 on the SK-BR-3 (SK) cells and PD-L1 in Alexa 488 on the EO771 (EO) cells. Differential interference contrast (DIC) images are shown on the left. F) Mean fluorescent intensity ratios (MFIR) were generated by normalizing each brightness value with the background and the average brightness of an untargeted cell. The HER2 and PD-L1 targeting was compared to the targeting of IgG controls. *n* = 8 – 15. *: *p*-value < 0.05.

### 2.2 Ultrasound imaging of mAb-GV conjugated by NHS and Staudinger ligation methods

With the cell characteristics tested and the mAb targeting confirmed optically, the mAb-GV conjugates could now be formed, targeted to cancer cells, and imaged with ultrasound. However, to the best of our knowledge a bioorthogonal site specific mAb-GV click conjugate had never been made to guide our next steps. Therefore, for this new protein-protein combination, we determined to use NHS-azide and NHS-phosphine reagents. After preparing HER2 and PD-L1 mAb-GV complexes using the NHS conjugation method and Staudinger ligation click method (Figure S1A,B,D), the cells were targeted for 90 min and then the unbound were washed away. The mAb-GVs bound to cells were loaded into a gel phantom for ultrasound imaging.

GV locations were visualized using difference images by subtracting images with cavitated GVs from images with intact GVs (Figure S2). In each image slice, the left sample contains untargeted cells (CTL), the middle contains cells targeted with the PD-L1 mAb-GV (PD-L1-GV), and the right contains cells targeted with the HER2 mAb-GV conjugate (HER2-GV). The EO771 results also reflect the preliminary tests (Figure 1A,C,D) with a high PD-L1 targeting and a non-significant HER2 targeting (**Figure 2**B). However, the SK-BR-3 cell results were not significant despite the high expression seen in preliminary tests (Figure 2A). The spleen and tumor cell results correlate to the primary cell flow cytometry (Figure 1B) with significant results for PD-L1 in both and HER2 just in the tumor (Figure 2C,D). This led us to hypothesize that the random binding of linkers and GVs on the mAb by the NHS method was possibly inhibiting the targeted efficiency.

**Figure 2.**
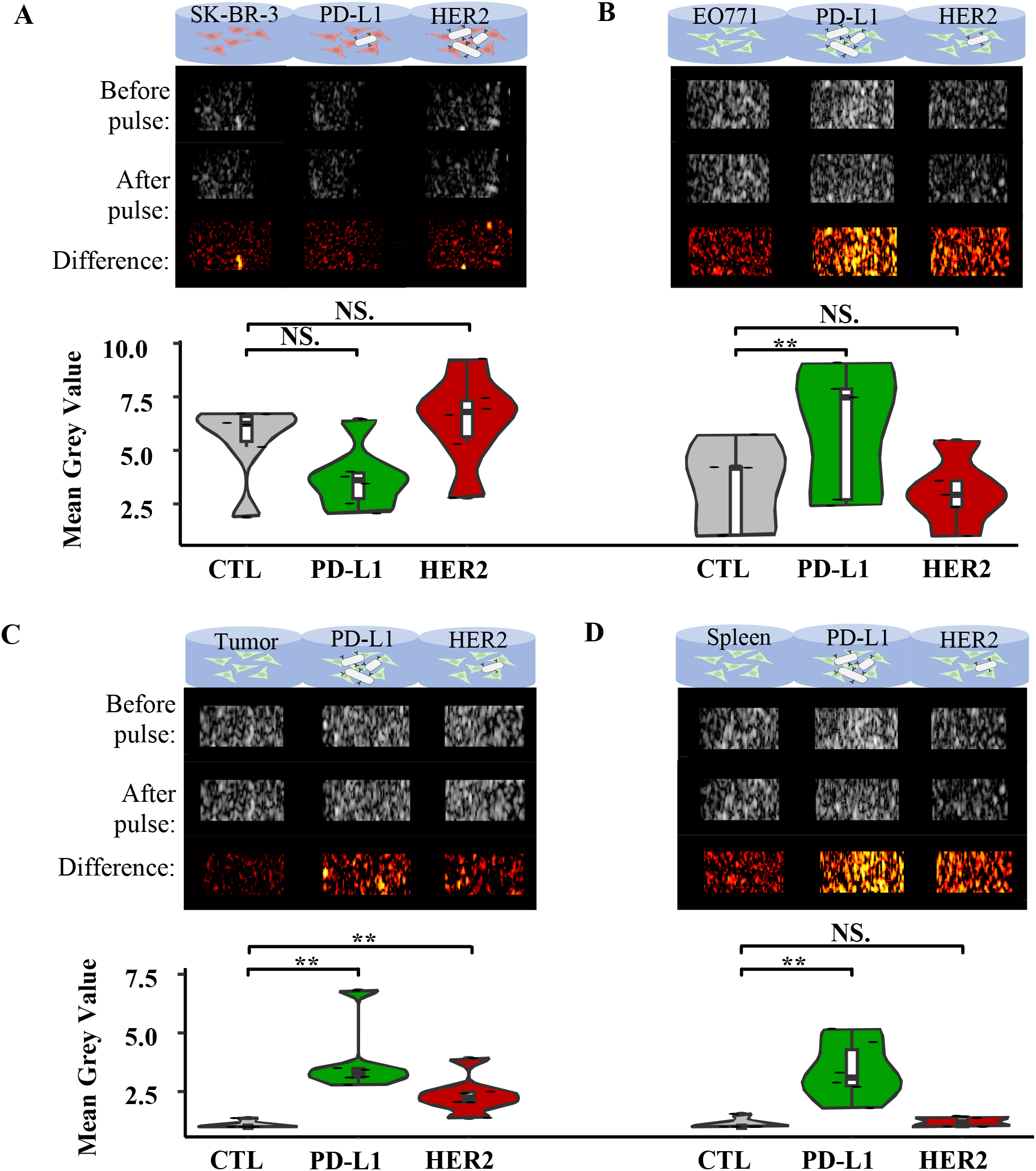
NHS mAb-GV ultrasound imaging. Ultrasound gel images were taken after targeting cells with HER2 and PD-L1 mAb-GV complexes with the NHS-azide conjugate and Staudinger ligation click chemistry (Figure S1A,B,D). For each of the four groups: the first column of well images are untargeted cells (CTL), the second column is cells targeted with PD-L1-GV, and the third is cells targeted with HER2-GV. A) SK-BR-3 results were not significant. B) EO771 results reflect the preliminary cell results from figure 2 with significance for the PD-L1-GV and not the HER2-GV. C) Targeting results using the primary cells from the tumors specifically matches the flow cytometry results in figure 1, with both PD-L1-GV and HER2-GV showing significance. D) Similarly, spleen cell targeting reflects the flow cytometry results with PD-L1-GV significance. *N* = 5 – 6. NS (not significant) > 0.05.^**^: *p*-value < 0.01.

### 2.3 Validation of Trop-2 targeting efficiency

To further validate the applicability of our method, we incorporated the targeting of Trop-2 by adding both a research mAb and a mAb used in clinic, Sacit. The MCF7 and HCC1806 targeted with anti-Trop-2 both showed significantly higher expression than the 293T control in the flow cytometry results (**Figure 3**A). Both Western blot and immunofluorescent tests also confirm high Trop-2 expression (Figure 3B,C). The research mAb and Sacit were each tested on both Trop-2 cell lines after fluorescent labeling (Trop-2-647) using the site-specific conjugation method, same as the HER2-647 and PD-L1-488 from figure 2E, showing significance for each (Figure 3D,E).

**Figure 3.**
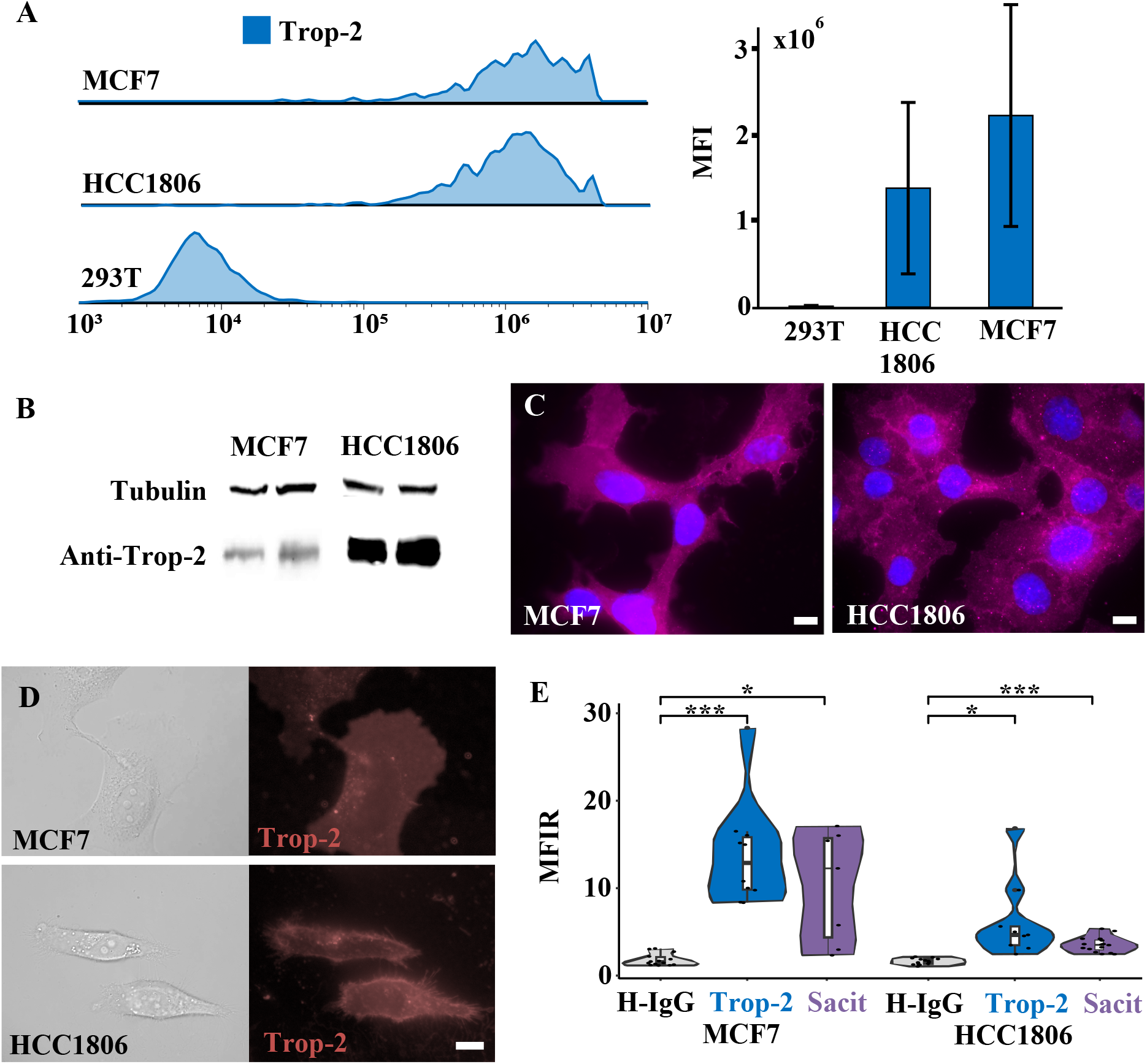
Trop-2 cell expression and antibody targeting efficiency. A) Flow cytometry, B) Western blot, C) IF imaging shows high expression of Trop-2 in MCF7 and HCC1806 cells. 293T cells were used as control. Blue indicates cell nucleus. Scale bar = 10µm. D) Site-specifically conjugated Trop-2-647 was used to target MCF7 and HCC1806 cells, showing high Trop-2 expressing in red. Scale bar = 10 µm. E) MFIRs were generated by normalizing each brightness value with the background and the average brightness of an untargeted cell. The Trop-2 targeting, both research mAb (Trop-2) and clinical drug Sacituzumab (Sacit), were compared to the targeting of a human IgG control. *n* = 6 - 14. NS (not significant) > 0.05. *: *p*-value < 0.05.^**^: *p*-value < 0.01. ^***^: *p*-value < 0.001.

### 2.4 NHS and site-specific GalNAz conjugation method comparisons

To overcome the limitations from the NHS method seen by the lack of SK-BR-3 targeting significance (Figure 2A), the site-specific GalNAz method was chosen due to the previous success for mAb fluorescent labeling. This method offers precise labeling to the glycans of the IgG heavy chain, enabling controlled and reproduceable conjugations. With both the NHS and site-specific conjugation methods prepared for the three mAbs, part was clicked to GVs and the other was clicked to fluorescent labels. The six mAb-GV samples were tested using a Zetasizer to determine their hydrodynamic range and zeta potential stability over time (**Figure 4**A). While there is an increase in size between the site-specific GalNAz and the NHS methods and after 2 weeks, the most significant increase occurs with the PD-L1 mAb, which also has the least stable zeta potential. The six mAb-647 samples were tested to measure protein concentration and fluorescent brightness to determine the degree of labeling (Figure 4B). As we hypothesized, the NHS method has many more labels due to the increased number of conjugation sites, but if those additions are on the binding region of the light chain, they may inhibit the cell targeting abilities of the mAb.^[18]^ Therefore, the two HER2 mAb-647 conjugates were used to mark SK-BR-3 cells to compare targeting abilities (Figure 4C). As expected, the cells targeted with mAb-647 conjugates made using the NHS method had higher fluorescent intensity due to increased Alexa 647 labeling, but the number of cells labeled by mAb-647 conjugated by the site-specific GalNAz method was higher.

**Figure 4.**
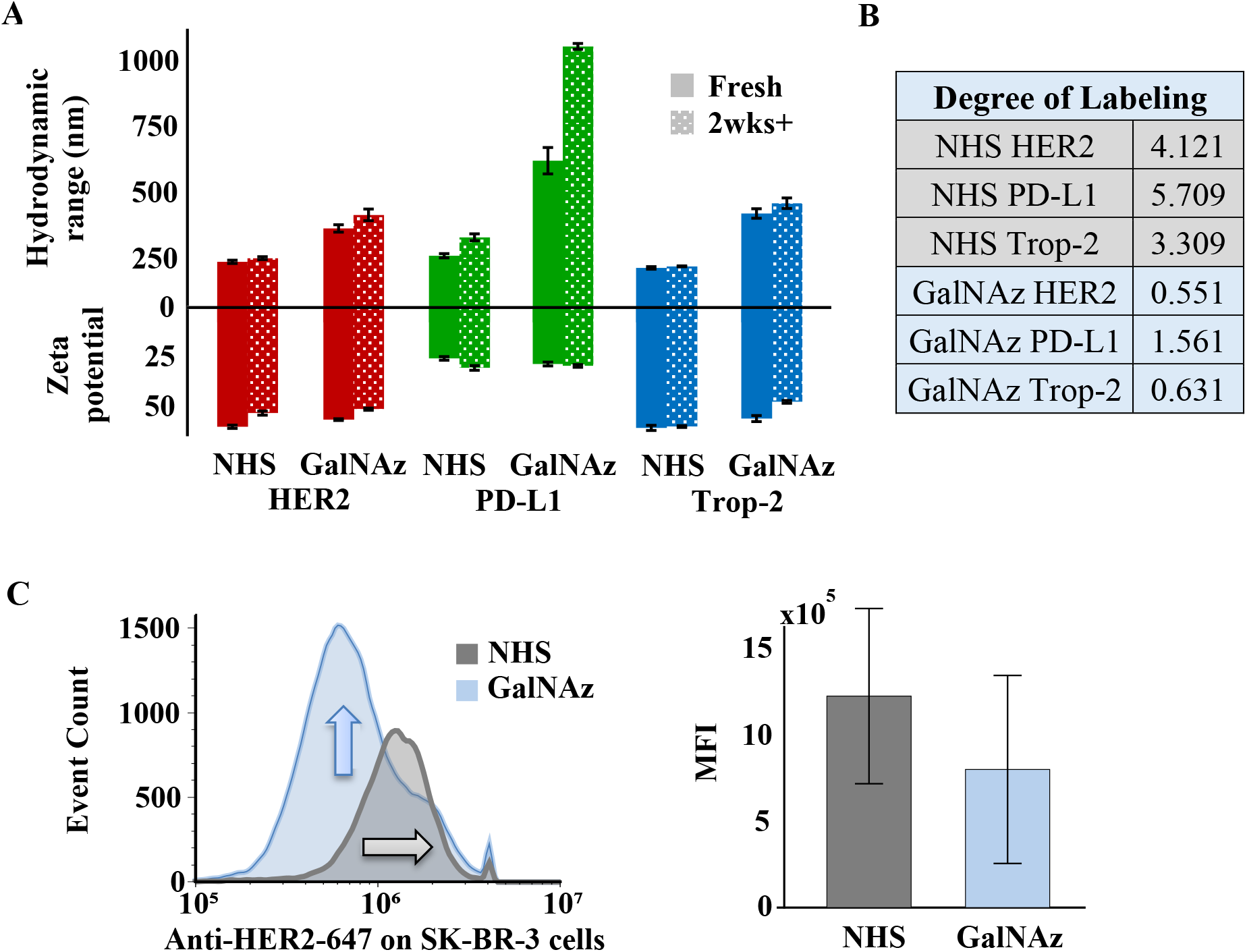
Conjugation comparison tests between NHS and site-specific GalNAz. A) Dynamic light scattering (DLS) measurement results using Zetasizer with the hydrodynamic range shown in the positive y-axis shows stability of conjugation between mAb and GVs. Zeta potential in the negative y-axis shows PD-L1 to be less stable, but not a significant difference between the NHS and site-specific methods. There is a slight decrease in stability for HER2-GV and Trop-2-GV. B) Degree of fluorescent labeling (DOL) for HER2, PD-L1, and Trop-2 with the NHS and site-specific methods using Alexa fluor 647 on the Nanodrop spectrophotometer shows a higher DOL for NHS verses site-specific and PD-L1 is higher than HER2 and Trop-2 regardless of the method. C) After targeted the same number of SK-BR-3 cells with HER2-647 conjugated either with the NHS or site-specific method, the flow cytometry results show NHS to be brighter (grey arrow), correlating to a higher DOL, but GalNAz to be more abundant (blue arrow), implying a higher targeted capability, which corresponds to the hypothesis of the mAb binding region being inhibited in the NHS method.

### 2.5 Ultrasound imaging of HER2, PD-L1, Trop-2 using site-specifically conjugated mAb-GV

After conjugation method comparisons showed both to be stable, the NHS method to have higher labeling, and the site-specific GalNAz method to have higher targeting, a final comparison was done by imaging targeted cells with ultrasound. As before, each image taken included three wells with untargeted cells on the left, the complimentary mAb-GV targeted cells in the middle, and the control IgG mAb-GV targeted cells on the right, but here each data point has been normalized using the brightness of the corresponding untargeted cells. Instead of comparing the brightness difference between the mAb-GV conjugate targeted cells and the untargeted cells as was done in figure 2, here we compared the brightness difference between the expected mAb-GV and the IgG-GV control, showing significance for all four cell lines using the site-specific conjugation method, but only two cell lines using the NHS conjugation method (**Figure 5**). This supports the hypothesis visualized in Figure S1D that the randomly binding NHS method may inhibit cell targeting and that site-specific binding may be preferred, especially for large additions such as GVs. Clinical drug Sacituzumab conjugates, Sacit-GV, targeted MCF7 and HCC1806 cells efficiently compared to the IgG control (Figure 5C,D). Unlike the fluorescent mAb conjugation, which used the commercially available Thermofisher SiteClick Antibody Azido Modification kit (click kit), we used site-specifically conjugated mAb-GV, which was developed and optimized in our laboratory according to the procedures described in the Experimental Section. For comparison, we also targeted SK-BR-3 and MCF7 with mAb-GVs conjugated using the click kit (Figure S3). The statistical significance shown for MCF7 was similar to that of Figure 5C and the SK-BR-3 significance decreases, suggesting the efficiency of our site-specific GalNAz method to develop ultrasound molecular contrast agent (mAb-GV)s.

**Figure 5.**
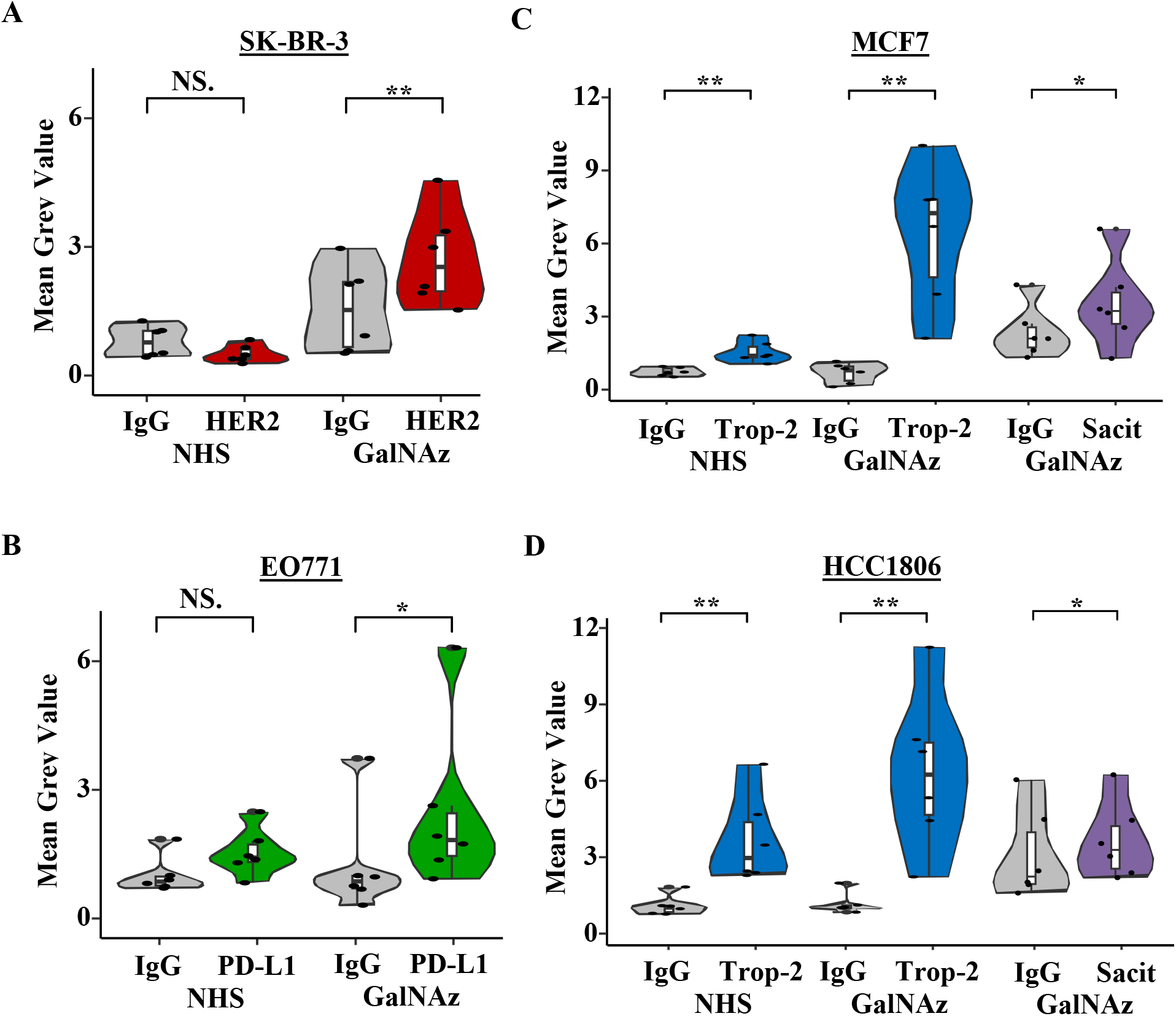
Ultrasound imaging comparisons between NHS and site-specific GalNaz. HER2, PD-L1, and Trop-2 mAb-GV complexes were prepared with the Staudinger ligation click method and either the NHS or site-specific GalNAz conjugation method to target the four cell lines and be imaged with ultrasound. As in figure 2, data is taken from the difference images. For each pair, the appropriate mAb-GV is on the right and the IgG control is on the left, and they have each been normalized with the brightness of a cell only well from the same image. For A) HER2 and B) PD-L1, the site-specific method is the only one that shows a significant increase from the control to the experimental. For Trop-2 targeting on C) MCF7 and D) HCC1806 cell lines, both the site-specific method and the NHS method are significant. Clinical drug conjugates, Sacit-GV, effectively target Trop-2 compared to IgG control. *n* = 6. NS (not significant) > 0.05. *: *p*-value < 0.05. ^**^: *p*-value < 0.01.

Though mAbs and GVs have each been studied *in vivo* separately, our site-specifically combined construct necessitated a biodistribution study in order to determine biocompatibility (Figure S4). 488-PD-L1 and 647-PD-L1-GVs were made by binding the fluorophores to the mAb carboxyl groups and adding the GVs using the site-specific method. The brain, lung, liver, spleen, and kidneys were tested for fluorescent brightness 1 hr and 2 hrs after i.v. injection of conjugates. The highest uptake of the 488-PD-L1 complex occurred in the kidney and liver, while the 647-PD-L1-GV was primarily found in the spleen and liver.

## 3. Discussion and Conclusions

Imaging cancer cells with enhanced ultrasound contrast by specific labeling of target biomarkers opens the door for a noninvasive diagnostic and monitoring option. The successful development of our bioorthogonal site-specific mAb-GV click conjugate forwards this end by providing a proven method for developing stable and precise probes that can be used for several cancer types. We demonstrated the limited targeting capability using the NHS method and the optimized conjugation with a site-specific method including the use of a mAb currently used in clinic. The GVs showed high stability because GVs are intact after 4 years (Figure S5) and the biodistribution test matched the expectations from literature, encouraging the next steps of testing.

The site-specific conjugation method significantly improved mAb-GV targeting of HER2 and PD-L1 compared to the NHS method. For Trop-2 targeting using MCF7 and HCC1806 cell lines, the statistical significance remained high for both methods. This inconsistency is likely due to the properties of each antibody, since the 10-15 lysine amine groups ready for NHS conjugation can be on different places for different mAb.^[18]^ Based on our results, it is possible that the Trop-2 mAb had few to no GVs conjugated to the binding region, since the NHS method also has a significant difference from the control. However, the brightness of the site-specific method may imply superior targeting regardless of similar p-value significance (Figure 5C,D).

Unlike the lysine amine groups, the site-specific method binding regions should not change from one IgG to another with two galactose regions on each glycan, making four targets that are each contained to the Fc region.^[19]^ This provides repeatability, predictability and unhindered cell targeting though it also decreases the labeling for additions like fluorophores (Figure 4B). For larger additions such as GVs, which are roughly 16 times longer (200 nm vs 12 nm) and 18 times wider (75 nm vs 4 nm) than mAbs, we would expect multiple mAbs on each GV, but no more than 1 or 2 GVs on each mAb due to lack of space. Therefore, the decreased DOL is less significant for the mAb-GV. Another factor for mAb binding is the stability. PD-L1-GV is less stable than HER2-GV or Trop-2-GV (Figure 4A) and has a lower site-specific ultrasound significance (Figure 5B). Conversely, PD-L1-GV has the highest DOL (Figure 4B) and the largest size increase (Figure 4A). Therefore, it is possible that while the instability may lead to increased conjugation, it may also lead to decreased targeting.

Regarding the biodistribution study (Figure S4), the complex made from conjugating Alexa 488 to the carboxyl groups of PD-L1 mAbs (488-PD-L1) (molecular weight (MW) of 153kDa and hydrodynamic range of 13 nm)^[49]^ was mainly cleared by the kidney, while the larger combination of Alexa 647 on PD-L1 mAbs site-specifically conjugated to GVs, (647-PD-L1-GV) was mainly cleared by the spleen and liver. If each 647-PD-L1-GV had one PD-L1 mAb and one GV, the MW would be approximately 190 kDa. However, multiple mAbs are likely to be conjugated to each GV, making the MW greater than 190 kDa. Based on this speculation, the calculated hydrodynamic range (Figure 4A) seems to suggest some clustering characteristics in the GalNAz PD-L1-GV complex, making it closer to 600 nm on average. Either way, the distribution was as expected since larger particles tend to be cleared by the spleen and liver, while smaller particles tend to be cleared by the kidneys.^[50]^

In summary, we conjugated mAbs and GVs using a bioorthogonal click reaction to develop ultrasound molecular contrast agents for cancer biomarker imaging. A precise and predictable conjugation with the specific targeting of multiple antibodies, including one currently being used in clinic, was achieved by utilizing the stability of GV and the site-specificity of the GalNAz method. Imaged cells targeted by developed mAb-GV conjugates demonstrated the molecular targeting and visualization of various cancer biomarkers. The proposed approach could supply the basis for a rapid and non-invasive cancer imaging and diagnostic tool that can be potentially used in clinic.

## 4. Experimental Section

### Gas vesicles production

AnaGVs were grown in BG-11 media (Thermofisher) from *Anabaena flos-aquae* (Culture Collection of Algae and Protozoa) for approximately 6 weeks in a CO_2_ controlled shaker with light cycles to mimic day and night. The GVs were lysed from the host bacteria using sorbitol and Solulyse, purified with PBS in a 4 day centrifugation buffer exchange process, and stored at 4°C at a concentration of 20 OD500.^[25, 51]^ For JEOL 2000FX Transmission Electron Microscopy (TEM) imaging, 2 μL of the GV solution was placed onto a carbon film copper grid (Electron Microscopy Sciences, EMS) and left for 3 minutes. A freshly centrifuges Uranyless solution of 5 μL was added to the grid and wicked after 30 seconds for imaging the next day.

### Primary cell isolation

C57BL/6 female mice were injected with 7.5 x 10^5^ EO771 cells from a 100 µl mixture with 50µl of cell solution and 50 µl of matrix gel in the fourth mammary fat pad and the tumor was allowed to grow until reaching 11 mm. After spleen and tumor dissociation and filtering, cells were targeted with a CD11b mAb (Biolegend), incubated, and washed. Magnetic nanobeads (Biolegend) were added to bind to the antibodies and the complex was filtered using a MAC separator and LSColumns (Miltenyi Biotec).

### Flow Cytometry

Cell solutions were mixed with a viability dye (Biolegend) before being targeted with fluorescently labeled mAbs (Biolegend) in a diluted serum. After imaging on a Cytek Northern Lights Flow Cytometer, populations were gated to eliminate debris, doubles, and dead cells. EO771 cells were activated with IFN-γ.

### Immunofluorescence imaging

Cells were fixed with paraformaldehyde (Sigma), permeated with Triton X-100 (Sigma), blocked with diluted serum, and targeted with a primary mAb (Cell Signaling) before being washed and targeted with RFP secondary mAb (Cell Signaling). DAPI was added to the cells to visualize the nucleus before being imaged using fluorescence microscopy.

### Western blot

Cells were lysed with RIPA buffer (Thermofisher) and loaded into protein gels (Biorad) before a 90-minute 70 V membrane transfer. The proteins were blocked with clear milk buffer (Thermofisher) for an hour and targeted with primary rabbit mAbs (Cell Signaling Technology) overnight at 4 °C on a rocker. After a wash, hour incubation with anti-rabbit secondary mAbs (Cell Signaling Technology), and another wash, chemiluminescence (Biorad) was used to image the membranes on a ChemiDoc.

### GalNAz Fluorescent Antibody Click

IgG mAbs (Bio X Cell) were labeled with azides using the SiteClick Antibody Azido Modification Kit (Thermofisher) and a prebound DIBO Alexa Fluor 647/488 (Thermofisher) was added before an overnight incubation and next day filtering. DOL alculation was used to determine the average number of successfully conjugated fluorophores. A NanoDrop spectrophotometer measured absorption (*A*), which were both used in conjunction with molar extinction coefficients (ε) and correction factors (*CF*) to calculate a ratio between the dye moles and the IgG moles (Equation 1).^[52]^

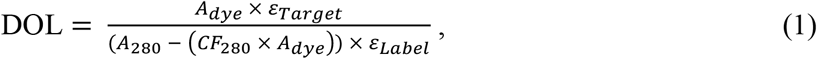

where A_dye_ and A_280_ are the absorbance readings given by the spectrophotometer. ε_Target_ for an IgG is a constant unitless 203,000, while the ε_Label_ for Alexa 488 and Alexa 647 are 73,000 and 234,000 respectively. The CF_280_ for Alexa 488 and Alexa 647 are 0.134 and 0.037 respectively.

### Antibody Targeting

Using two adherent wells (Ibidi) on a glass bottom dish, the cells were added so that the control and the experimental group were in the same dish. The appropriate fluorescently conjugated mAbs were added, using conjugated human IgG controls for the human cell lines and a mouse IgG control for EO771, and incubated for 90 minutes. The extra mAbs were washed away, and the cells were imaged using fluorescent microscopy.

### NHS mAb-GV Click

Both NHS ends of the NHS polyethylene glycol 4 (PEG4) azide (Vector Laboratories) and NHS phosphine (Thermofisher) click ingredients were prepared using dimethyl sulfoxide (DMSO) and then added to the target proteins to produce azide-mAb and phosphine-GV. After an hour at room temperature, the reaction was neutralized using a tris quenching buffer. To remove excess chemicals, the azide-mAb was filtered through a molecular weight cut off (MWCO) tube and the GV-phosphine was washed with two rounds of centrifugation and buffer exchange. The two prepared proteins were then combined with a mAb molar excess and incubated overnight in 4°C. A final wash was done using the centrifugation and buffer exchange process.

### GalNAz mAb-GV Click

After 1 mg of the mAb had been buffer exchanged into sodium acetate (Sigma) using MWCO tubes, 20 µl of *β*-galactosidase and 20 µl of the corresponding buffer (New England Biolabs) were added and incubated overnight at 37 °C to remove the galactose molecules from the mAb. The next day, 440 µg of UDP-N-azidoacetylgalactosamine disodium (UDP-GalNAz), 50 µg of β-1,4-galactosyltransferase (GalT) (MedChemExpress), 18 µl of 20X tris(hydroxymethyl)aminomethane (Tris) (Sigma), 10 µl of 10X Tris-buffered saline (TBS) and 12 µl of Type 1 water were added for another overnight incubation, but this time at 30 °C. The excess chemicals were washed away using MWCO columns and the site-specifically conjugated azide mAb was added to NHS-phosphine prepared and washed GVs with a mAb 2X molar excess for a third overnight incubation, now at 25 °C. After the final wash, the mAb-GV conjugates were stored in 4°C.

### mAb-GV cell targeting

For targeting adherent cells, three 35 mm dishes were used for each cell line; 1 for untargeted cells and 2 for targeted cells. Twelve 70 µl volume well inserts (Ibidi) were used for each of the targeted cell dishes and an equal number of cells were plated in the dish for untargeted cells. Since GVs float, the dish was turned upside down, keeping the liquid inside via well wall surface tension, and allowing the GVs to float up to the cells for 90 minutes of binding. The suspended primary cells were also targeted for 90 minutes, but via a rotator in the incubator.

### Gel Phantom

Cylindrical gel wells were made in a plastic container using a 3D printed lid with 24 small, raised columns in 8 rows of 3 sized to fit in the frame of the linear array transducer. A 1% agarose (Biorad) gel was made by heating 1 g for agarose powder in 100 mL of Type 1 water. After pouring the liquid gel into a plastic container, the lid was placed on top, and the solution solidified at 4 °C for 10 minutes. When the lid was removed, gel well sample holders were left imprinted in the larger gel area. The targeted and pelleted cells were resuspended in media to make two identical samples of 5 μL for each type. Quickly 20 μL of gel was added to the cell mixture, swirled, and loaded into a gel well. The gel container was then stuck down into a larger container, covered with water, and imaged using a linear array transducer.

### Ultrasound imaging system

Ultrasound signal transmissions and corresponding radiofrequency (RF) and in-phase and quadrature (IQ) data acquisitions were performed using the Verasonics Vantage 256 ultrasound research system (Verasonics, Seattle, WA) with an L22-14vX linear array transducer (15.6 MHz center frequency, 128 elements). The transducer was mounted on a motorized 3D translation stage (ILS150CC, Newport Corp., Irvine, CA) for accurate positioning and mechanical stability throughout imaging. All data processing was performed in Matlab on a desktop computer with an Intel® Xeon® W-2155 3.3 GHz processor and 32 GB RAM._[53]_

### Pulse sequence and image acquisition

The gel-well phantom was first imaged using a 1.5-cycle sinusoidal pulse having a pulse width of 20 ns with a peak-to-peak voltage (V_pp_) of 10 V, to obtain pre-cavitation (baseline) image. Cavitation was then induced by transmitting 1.5-cycle pulses with V_pp_ of 30 V, having the same pulse width, at a pulse repetition frequency (PRF) of 500 Hz for 1 second to cavitate the GVs. After cavitation, the phantom was imaged again with the same parameter of pulse as the pre-cavitation image, to obtain the post-cavitation image. The phantom was kept firmly stationary throughout the entire process to prevent motion artifacts.

### Difference image processing

The IQ data of the pre and post cavitation frames were normalized within the same dynamic range and the post-cavitation frame was directly subtracted (pixel-by-pixel subtraction) from the pre-cavitation frame, and only positive changes were retained. This process generated the difference image that captured the changes that took place in each well during the cavitation.

### Biodistribution study

The ethylene dichloride (EDC) substitution reaction was used to bind Alexa fluor 488 carboxylic acid (Thermofisher) to the PD-L1 mAb (Bio X Cell) carboxyl groups, to produce 488-PD-L1. A second PD-L1 mAb complex was made using the EDC reaction, but this time using 647 and with the addition of GVs using the site-specific GalNAz method to make 647-PD-L1-GVs. Both complexes had the fluorophores washed away using the snakeskin dialysis method in PBS and after the GV addition on 647-PD-L1 an additional wash was preformed using the centrifugation buffer exchange. The GVs had been prepared and washed similarly to the NHS phosphine addition, but this time with NHS PEG4 dibenzocyclooctyne (DBCO). Six C57BL/6 female mice were injected with 488-PD-L1 and six with 647-PD-L1-GV to test 3 of each after 1 hr and 3 of each after 2hrs. The brain, lung, liver, spleen, and kidneys were weighed, dissociated, and imaged on a plate reader for fluorescent brightness, normalized by organ weight.

### Statistical Analysis

P-value significance tests were performed using Excel and RStudio, using a paired addition for the ultrasound results to focus the comparison to the same image slice.

## Acknowledgements

This work was supported in part by the National Institute of Health (NIH) grant Nos. GM120493 and GM152704 (to S.Y.), the National Science Foundation (NSF) grant Nos. CBET1943852 and 2304485 (to S.Y.), NIH R01 CA255064-01A1 (to S.Z.), and the Cancer Prevention and Research Institute of Texas Scholar Award RR220024 (to S.Z.). Figure 1B was created using Biorender.com. ToC figure was created with images from pdb101.rcsb.org and Biorender.com

## Data Availability Statement

The data that support the findings of this study are available from the corresponding author upon reasonable request.

